# Exploiting facial side similarities to improve AI-driven sea turtle photo-identification systems

**DOI:** 10.1101/2024.09.13.612839

**Authors:** Lukáš Adam, Kostas Papafitsoros, Claire Jean, ALan F. Rees

## Abstract

Animal Photo-identification (photo-ID) denotes the process of identifying individual animals based on their unique and stable morphological characteristics. It has been proven to be a particularly useful tool for a variety of studies on sea turtles increasing the knowledge of their ecology and informing conservation efforts. The photo-ID process in sea turtles is predominantly based on the geometric patterns of the scales of their two head sides which are unique to every individual, and also different from side to side. As such, both manual and automated photo-ID techniques are performed under a side-specific image retrieval setting. There, a query image showing a single profile of an unknown individual, either left or right, is only compared to images of previously identified individuals showing the same side. Here, by employing the recently introduced state-of-the-art deep feature extractor MegaDescriptor, we show for the first time an inherit deep visual similarity between left and right facial profiles of the same individuals in three sea turtle species. We show that the similarity between the left and right profiles of the same individual with respect to geometry, coloration and pigmentation, is on average higher than the similarity between profiles of different individuals. The similarity is detectable even when images are taken years apart and under diverse settings and conditions. We perform several image retrieval experiments under scenarios which mimic realistic sea turtle photo-ID matching processes, where we also allow comparisons of opposite sides in the matching process which have no spatial overlap. We show that the detection and exploitation of this similarity is translated to improved accuracies when compared to the traditional side-specific image retrieval setting. Notably this similarity cannot be detected and thus neither explored by the so-far state-of-the-art sea turtle photo-ID automated methods such as those based on SIFT. Our work leads to a change of paradigm for the sea turtle photo-ID workflows which will be further facilitated by the constant improvement of deep feature-based re-identification methods and paves the path for adopting similar workflows in other animal species as well.

## 1. Introduction

Animal photo-identification (photo-ID; commonly refereed to as *re-identification* in the computer vision literature) denotes the task of identifying individual animals from images by exploiting their unique and stable in time external morphological patterns. Photo-ID is particularly widespread in studies of wildlife populations of various taxa and species like cetaceans (Cheeseman et al., 2017), sharks (Araujo *et al*., 2017), sea turtles (Schofield *et al*., 2020), rays (Marshall and Pierce, 2012), bears (Anderson et al., 2010), seals (Koivuniemi et al., 2016) giraffes & zebras (Parham et al., 2017) to name a few representative works. It overcomes logistical difficulties of capturing and tagging the animals using external tags, while being minimally invasive causing minimal or no stress and it can be performed in large scales when it is combined with active or passive citizen science (Holmberg et al., 2009; Papafitsoros et al., 2021).

Sea turtles were one of the first group of species to which researchers applied photo-ID (McDonald and Dutton, 1996) and its use was further intensified after the popularisation of digital cameras (Schofield *et al*., 2008). All seven sea turtle species are keynote species, globally protected under international laws whose survival is crucially relying on continuous research and conservation efforts (Rees et al., 2016). Today, photo-ID is used as an indispensable tool in a variety of sea turtle studies like survival rate estimation (Schofield *et al*., 2020), in-water behavioural studies (Chassagneux *et al*., 2013; Schofield *et al*., 2017; Schofield *et al*., 2022), injury and disease monitoring (Bennett *et al*., 2000; Ciccione *et al*., 2015; Papafitsoros *et al*., 2021; Hancock *et al*., 2023), measuring ecotouristic pressure (Hayes *et al*., 2017; Papafitsoros *et al*., 2023), inferring population distribution (Hanna *et al*., 2021; Hudgins *et al*., 2023; Neves-Ferreira *et al*., 2023), see also Papafitsoros *et al*., 2024 for a comprehensive review.

A major challenge in sea turtle, and generally animal photo-ID is that it can often be time consuming due to the ever increasing number of obtained images. Speeding-up the process is possible by employing divide-and-conquer strategies, e.g. by splitting photo-databases into sub-databases of much smaller size, each one corresponding to a particular characteristic e.g. location, sex or a fine categorisation of the identifying patterns (Schofield et al., 2008; Lloyd et al., 2012; Papafitsoros et al., 2024). However, this only partially solves the scalability problem and it is not always applicable. This need to reduce manual labour has been driving the development of fast and accurate automated photo-ID methods which we discuss in detail in Section 3.

Sea turtle photo-ID is predominantly based on the animal’s facial scales polygonal pattern which is a unique to every individual. It has been shown that these patterns are stable in time throughout the whole animals’ lives (Carpentier et al., 2016). With the exception of using drones for sea turtle photo-ID (Comis et al., 2022) where the top side of the head is used, the identification is mainly based on the left and right sides of the head (Papafitsoros *et al*., 2024). Both manual and automated sea turtle photo-ID are “classically” performed under a side-specific image retrieval setting: Any new image (*query image*) depicting a side of the head of an unknown individual, is visually compared to a group of images (the *database*) which consists of images of the same side of previously known individuals, until a *matching* occurs, see Figure 1(a). The left and right polygonal scale patterns are different in a given individual, and thus sea turtle photo-ID workflows that include comparisons of opposite sides, see Figure 1(a), currently do not exist. As a result, if only the opposite side to the one of the query image is available in the database, the individual cannot be identified. This is a common problem due to e.g. incomplete submissions by citizen scientists or skittish animals. However, it has recently been reported (Papafitsoros et al., 2024) that despite the different patterns, there is still some inherit visual similarity between left and right profiles at a given individual, not only in coloration and texture but also geometrically, see Figure 2 in the next section. Yet, this has so far neither been quantified nor been exploited in sea turtle photo-ID.

**Figure 1.**
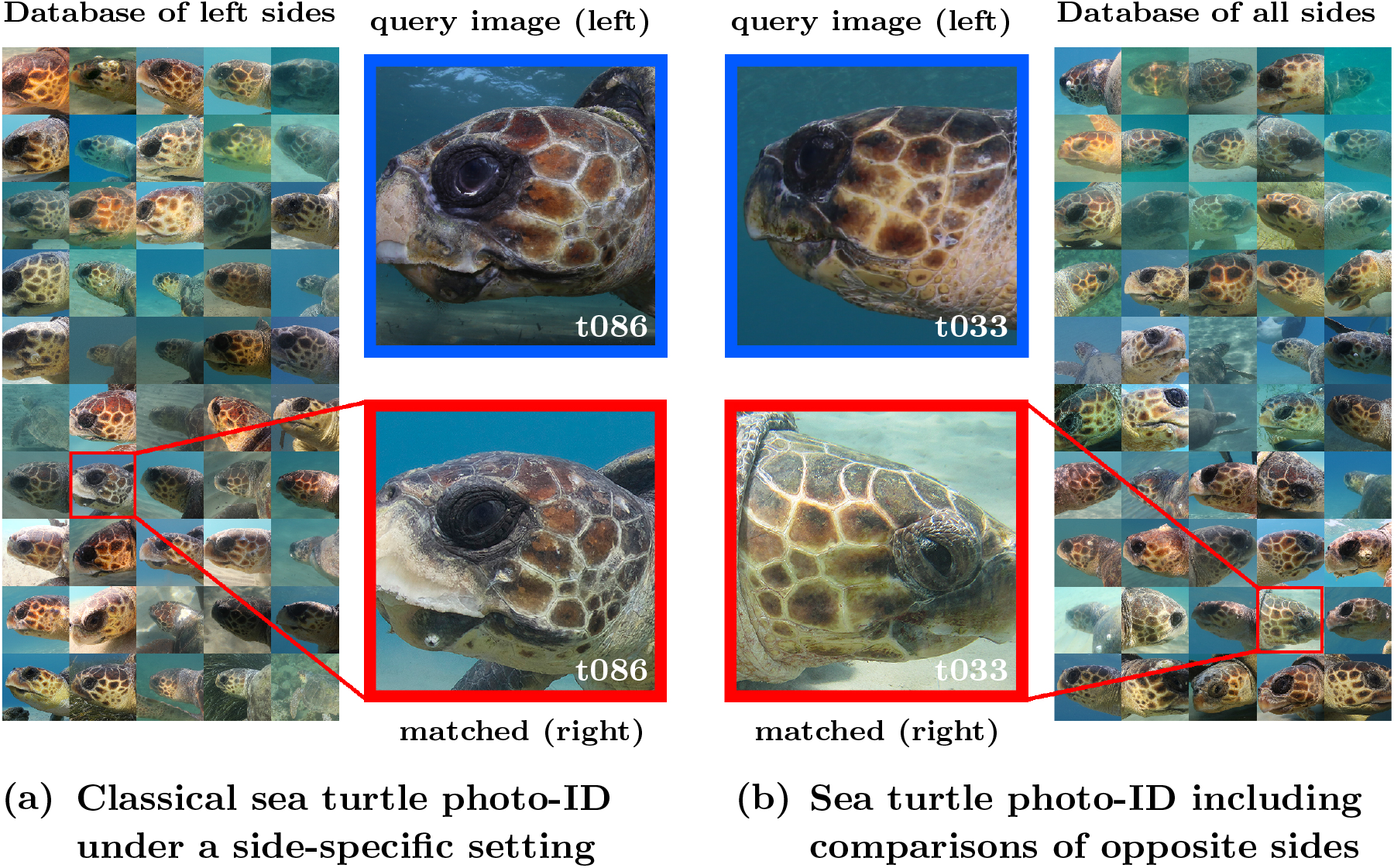
(a) “Classical” sea turtle photo-ID workflow where a query image (here left side of the head) is compared to a database of images of the same side (here only left sides). The eventual matching is based on the exact similarity of the polygonal patterns which remain stable across time. (b) Workflow where also comparisons of opposite sides are performed. So far, manual and automatic photo-ID methods have been following the first approach. In this work, we show that the second approach is also feasible and beneficial when a deep feature re-identification method like MegaDescriptor is used. In this schematic, the image showing left side of the individual “t033” is matched to an image showing its right side, since this is the most similar one in the database. Note that the left side of “t033” is not present in the database. All images are taken from the SeaTurtleID2022 dataset (Adam et al., 2024a)

**Figure 2.**
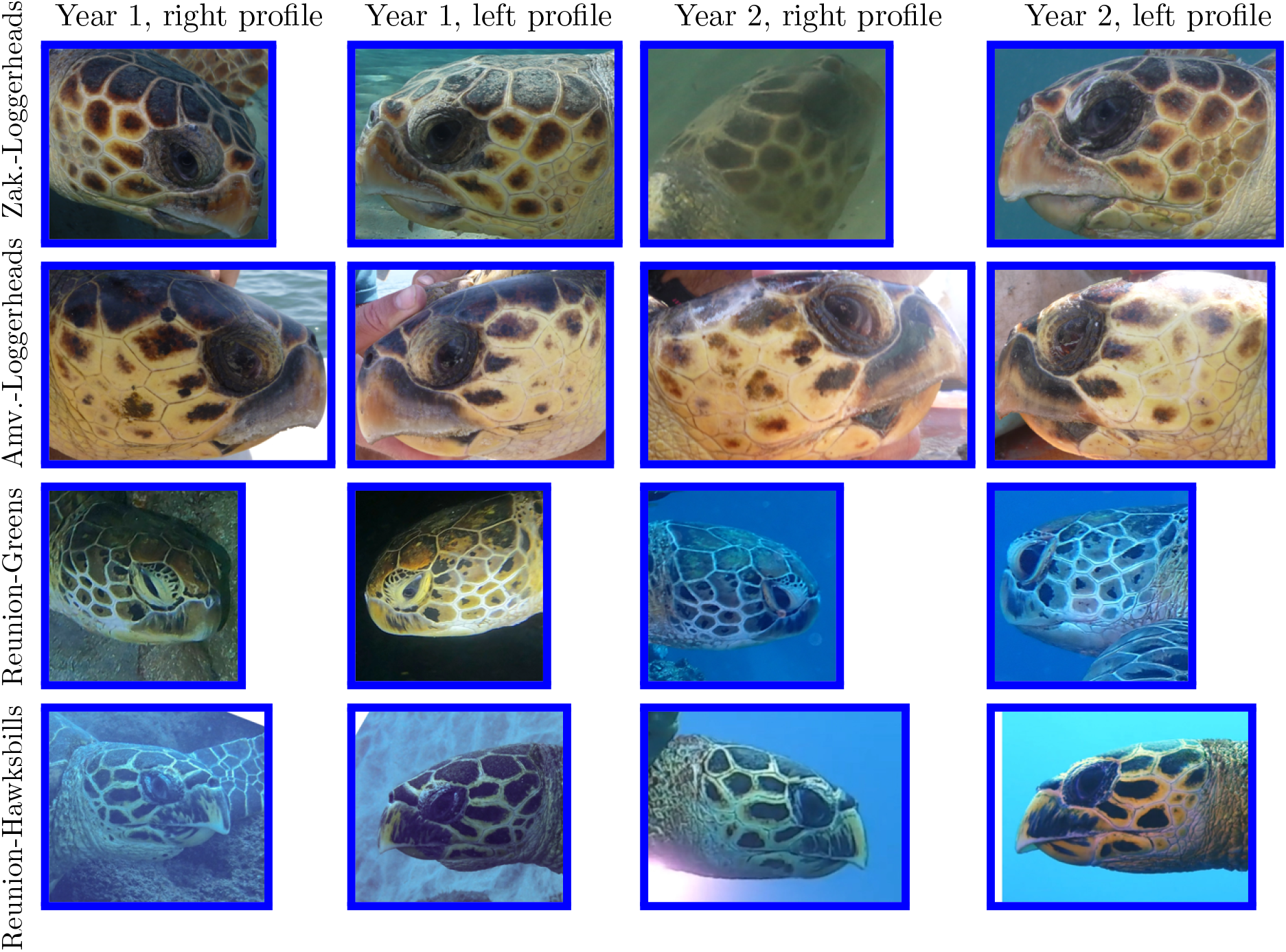
Sample photos for individuals (one single individual per row) belonging to each of the 4 considered datasets. From every individual four photos were chosen in total, taken in two different years with both profiles being represented in every year.

**Our contribution** In this work, we make the following contributions:

i. We show that a recently introduced deep feature-based model for wildlife re-identification, MegaDescriptor (Čermák *et al*., 2024), can be successfully used for sea turtle photo-ID in a variety of photo-capturing settings and species. We test it on 4 multi-year spanned databases of the three common sea turtle species for which photo-ID is employed: Loggerhead (*Caretta caretta*; two settings), Green (*Chelonia mydas*) and Hawksbill (*Eretmochelys imbricata*) sea turtles.
ii. By employing MegaDescriptor’s capability to produce meaningful similarity scores for pairs of animal images, we show for the first time in a quantitative way, the inherit *deep* visual similarity between left and right facial profiles of the same individual in these three sea turtle species, which so far has only be described empirically. In particular, we show that the similarity between the left and right profiles of the same individual is on average higher than the similarity between profiles of different individuals. This similarity is more prominent for images taken in the same year but it is detectable even when images are taken years apart and under different settings/conditions. In contrast we show that local feature methods (SIFT) cannot detect and thus cannot exploit this similarity.
iii. We show that opposite profiles of an individual taken in the same year are ranked more similar than same profiles of the same individual taken in different years, suggesting changes over time in coloration, pigmentation and other subtle non-geometrical features. On the other hand, when comparing images taken in different years, the choice of sides plays a not as significant role as previously thought. This means that having only a single profile of an individual in a database, poses no significant restrictions for automated identification.
iv. We perform a series image retrieval experiments under a variery of scenarios which mimic realistic sea turtle photo-ID matching processes, where we also allow comparisons of opposite sides in the matching process which have no spatial overlap. We show that the detection and exploitation of this similarity is translated to improved accuracies when compared to the traditional side-specific image retrieval setting.
v. Finally, in order to encourage and to standardise further research in this novel direction of sea turtle photo-ID we make our datasets and codes publicly available (links provided at the end of the paper).

## 2. Testing datasets

Before we proceed in describing our methodology and experimental design, we present first the different datasets used to perform our experiments. We used four distinct datasets, two of loggerhead sea turtles (one of underwater photographs and one of out-of-water ones), as well as one of green and one of hawksbill sea turtles, both of underwater photographs. For all images in all datasets, the following metadata were available: turtle identity, head orientation (left or right) and timestamp (year that each photograph was taken). We briefly describe next the characteristics of each of these four datasets.

### 2.1. Dataset Zakynthos-Loggerheads

This dataset consists of photographs taken between 2018 and 2024, in Laganas Bay, Zakynthos Island, Greece (37^°^43^*’*^N, 20^°^52^*’*^E), which is a main breeding site for the Mediterranean loggerhead sea turtles (Margaritoulis et al., 2022). All photographs were captured underwater by the same photographer (KP) during snorkeling surveys from a distance ranging from 7 meters to a few centimeters. A Canon 6D full-frame DSLR camera (5472×3648 pixels) combined with a Sigma 15mm fisheye lenses was mainly used. A small number of the photographs were captured using a Canon R8 mirrorless camera with either the Sigma 15mm or a the Tokina 10-17mm lenses. The water depth ranged from 1 to 8 meters, with the vast majority of photographs taken less than 5 meters deep.

This dataset consists of images of the same *distribution*, i.e. same capture conditions and sea turtle population, as the SeaTurtleID2022 dataset (Adam et al., 2024a) which was part of the training set of the MegaDescriptor, see Section 3.2. However, it consists of new individual loggerheads which were not included in SeaTurtleID2022 and thus were not seen during the training of MegaDescriptor.

### 2.2. Dataset Amvrakikos-Loggerheads

This dataset also consists of photographs of Mediterranean loggerhead sea turtles taken at Amvrakikos Gulf, Greece (39^°^02^*’*^N, 21^°^06^*’*^E) which is wellknown foraging site for adult and juvenile turtles. Photographs were collected as part of a long-term capture-mark-recapture project conducted by ARCHELON, the Sea Turtle Protection Society of Greece (Rees et al., 2013). Turtles were captured from a boat using the sea turtle rodeo technique and among other data collected, photographs of the head sides were taken while the animal was on the boat. In all photographs, either the whole side of the head was fully shaded or fully illuminated by the sun. All photographs in this dataset were taken during the summer months (June-August) between 2014 and 2022, using a selection of different digital cameras of varying optical resolution. While it is known that many of the turtles foraging in Amvrakikos Gulf are genetically similar with the ones of Zakynthos (Rees et al., 2017) and thus share common morphological characteristics, the different conditions of photograph collection make it a distinct dataset to the Zakynthos-Loggerheads one.

### 2.3. Datasets Reunion-Greens & Reunion-Hawksbills

This dataset consists of photographs taken between 2007 and 2024 on Reunion Island, a French territory in the Indian Ocean (21^°^06^*’*^S, 55^°^30^*’*^E), whose shallow waters are known to be a development and foraging ground for green and hawksbill turtles (Chassagneux *et al*., 2013). Photographs were taken by recreational divers as part of a citizen science programme with no specific associated protocols (no specific viewing angle or distance). Citizens shared their photographs whenever they wished with scientists for further analysis. All photographs are turtle head profiles taken during the same encounter and are stored in the TORSOOI database, see Section 3.4 for more details.

The links to all four datasets can be found at the end of this paper.

### 2.4. Data selection and preprocessing

We selected 40, 50, 50 and 34 individual turtles for Zakynthos-Loggerheads, Amvrakikos-Loggerheads, Reunion-Greens and Reunion-Hawksbills, respectively. We note again that for Zakynthos-Loggerheads we used individuals not included in SeaTurtleID2022 (Adam et al., 2024a), which is a dataset containing images of 400 individuals from Zakynthos Island. We provide a short summary of the selected photos for each dataset in Table 1.

**Table 1.** Overview of the four testing datasets: Number of individuals, total number of images (number of individuals 4), conditions of photo-capturing and average time-span of photographs in years.

| Dataset | Individuals | Images | Conditions | Average Span |
| --- | --- | --- | --- | --- |
| Zakynthos-Loggerheads | 40 | 160 | underwater | 2.5 years |
| Amvrakikos-Loggerheads | 50 | 200 | on a boat | 4.4 years |
| Reunion-Greens | 50 | 200 | underwater | 4.7 years |
| Reunion-Hawksbills | 34 | 136 | underwater | 3.4 years |

For every individual in each dataset we used four photographs for our experiments: two (left and right side) taken during a given year and two (left and right side) taken during a different year, see Figure 2 for sample photos from each dataset.

The main reason for including images taken during different years is to eliminate the undesirable phenomenon when an algorithm decides that two images contain the same turtle, not based on the morphological patterns of the turtle but rather on some external factors such as similar background or lightning conditions. Images of the same individuals taken during different years are not expected to share the latter two and are only matched due to turtles’s morphological characteristics (Adam *et al*., 2024a). To further reduce any background effect, we applied a bounding box around each turtle’s head, that is, all the experiments were performed on the turtles’ heads only. On the other hand, images taken during the same year, especially the ones taken underwater during the same encounter might share the same *global* lighting conditions, e.g. see the images of “Year 2” at the third row of Figure 2. In order to account for this and examine its influence on the identification process, we also converted all images to grayscale and performed all the experiments twice, once for coloured and once for grayscale versions. We show the grayscale analogue of Figure 2 in Figure For instance, focusing on the right profiles of the turtle in the third row we notice that the differences in global coloration in Figure 2 are not present any more in the grayscale version in Figure 3. We stress however that the conversion to grayscale also suppresses the individual turtles’ skin coloration. This might prevent detecting its natural color change, which is known to occur over the years (Carpentier et al., 2016; Adam et al., 2024a; Papafitsoros et al., 2024), as well as the across-individual different skin coloration which apart from the geometrical patterns also informs the re-identification. We note that for images taken outside of water (Amvrakikos-Loggerheads; see second row of Figure 2), there are minimal global lighting differences across the years, in comparison to underwater photos.

**Figure 3.**
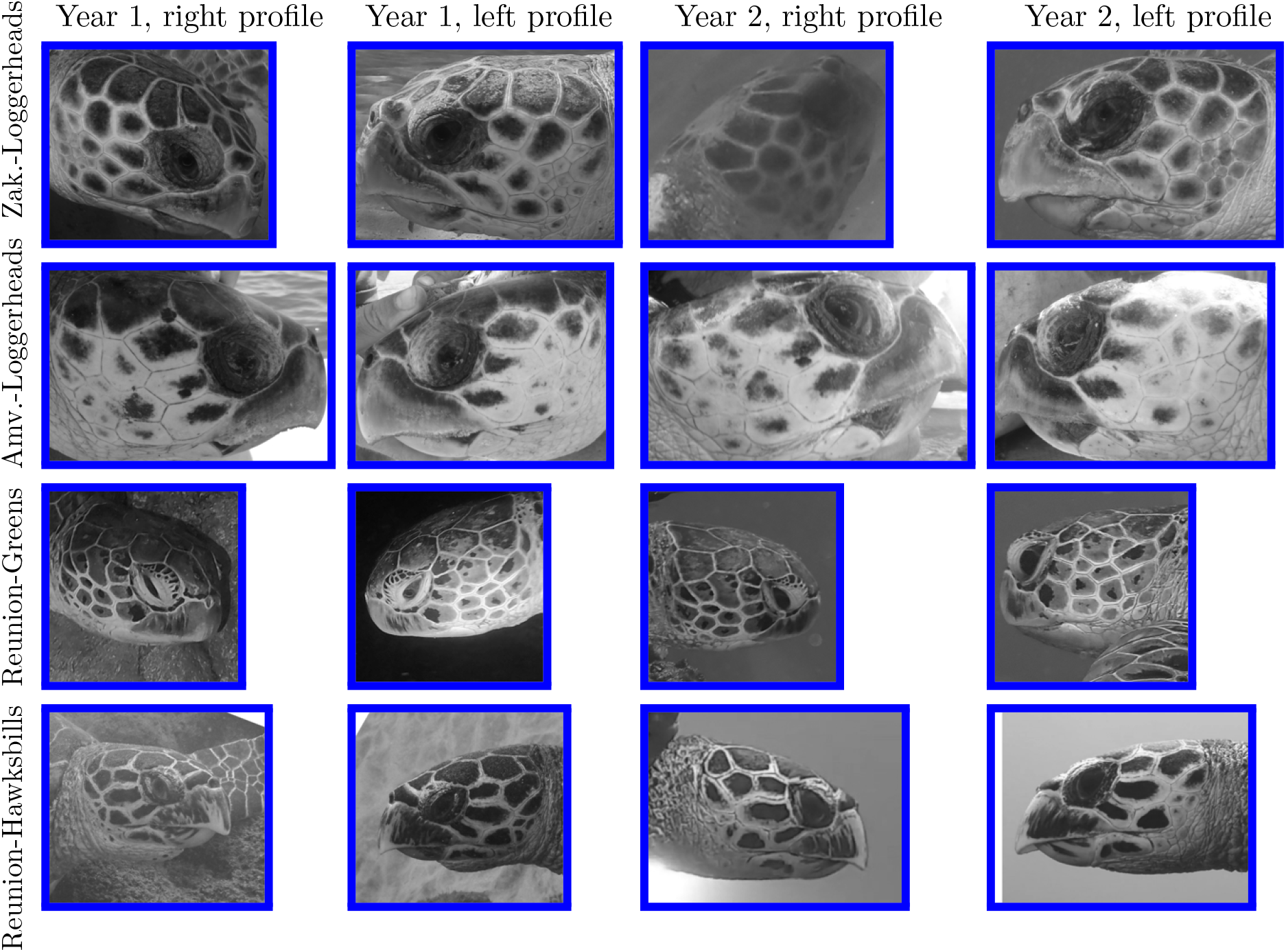
Grayscale version of Figure 2, eliminating global coloration differences due to different capture settings, e.g. compare the images of the individual in the third row of the two Figures. We also stress however that conversion to grayscale suppresses the natural skin coloration differences which can be found both among different individuals but also on same individuals over different years. Here we emphasize that grayscale conversion is only a part of the method evaluation process and should not be seen as part of the standard sea turtle photo-ID practice.

## 3. Methodology

This section describes the used algorithms and the experimental design. We first describe the general framework of image retrieval which is based on similarity comparisons between images and we give a short overview of automated methods that compute these similarities. We then proceed in describing in more detail three such methods used for our experiments: MegaDescriptor (a recently introduced state-of-the-art deep feature-method), SIFT (a classical local feature-based method) and TORSOOI (a sea turtle-specific method). We then describe our experimental design which includes several settings for examining the similarity of sea turtle head profiles, and a series of related image retrieval experiments and their evaluation.

### 3.1. Image retrieval and automated photo-ID methods

The majority of automated animal photo-ID methods operate under an image retrieval setting. Image retrieval is a standard technique in computer vision to evaluate classification algorithms. Having a single query image (to be identified) and a database (images of already identified individuals), it selects the “most similar” image from the database to the query image. The query image is then predicted to depict the same individual as the selected database image. The goal of an algorithm is to compute the similarity measure so that it is high for images with the same individual and low for images with different individuals. Since image retrieval retrieves an image (and not an identity), it allows for easy manual verification of the prediction.

Following Čermák *et al*., 2024, automated methods can be divided into three categories: (i) local feature-based, (ii) deep feature-based and (iii) species-specific methods. The three methods described later represent each one of these categories.

Local feature-based methods extract local descriptors (local image features of salient characteristics e.g. edges) from the query image and subsequently match them with the corresponding extracted descriptors of the database images. Prominent examples for these features are the keypoints extracted by the Scale Invariant Feature Transform (SIFT) (Lowe, 2004), see Section 3.3, or the Speeded Up Robust Features (SURF) (Bay *et al*., 2008). For instance, methods based on SIFT (e.g. Hotspotter (Crall *et al*., 2013); see also Dunbar *et al*., 2021 for its application to sea turtles) detect a series of keypoints, and rank the database images according to the degree that their keypoints match. Local feature-based methods typically require no training and fine-tuning but they face challenges in scaling efficiently to large datasets.

On the other hand, deep feature-based methods exploit the power and the expressivity of deep neural networks (Goodfellow et al., 2016) to represent an image of an individual via a *deep feature vector* which has a much lower dimension than the image. The neural network is trained on sufficiently large databases and *learns* how to efficiently encode characteristics of individual animals to these feature vectors (Čermák *et al*., 2024; Shinoda and Shiohara, 2024). These identifying characteristics can be richer than local features, and can include global geometric patterns, coloration, texture etc. In contrast to local-feature-based methods, these methods are largely benefitted when trained on groups of images that are similar to the ones that are later applied to, e.g. same or similar species. We note that the application of deep feature-based methods to sea turtle photo-ID remains relatively unexplored.

The species-specific methods exploit some particular morphological characteristics of the focal species and cannot be used for other species. Examples include (Anderson et al., 2010) and (Kelly, 2001) designed exclusively for polar bears and cheetahs, respectively. For sea turtles we mention TORSOOI (Jean et al., 2010) which encodes the facial polygonal pattern in a single numerical code and it is described in more detail in Section 3.4.

### 3.2. MegaDescriptor

is a recently introduced deep feature vector extractor (Čermák et al., 2024). It was trained in such a way so that images of the same (resp. different) individuals have similar (resp. distinct) deep feature vectors. MegaDescriptor is freely available from HuggingFace. It is written in Python and was developed putting emphasis on its simple use, e.g. the feature extraction is performed by one line of code. We briefly sketch how MegaDescriptor is used under an image retrieval setting in Figure 4. The left-hand side shows the query image and the right-hand side the database. A feature vector is extracted from each image, and the similarities between feature vectors are computed using the cosine similarity metric. Then the query image is predicted to show the same individual as the database image with the highest similarity. We note that in principle there is no restriction that the database image should depict the same side of the head as the query image.

**Figure 4.**
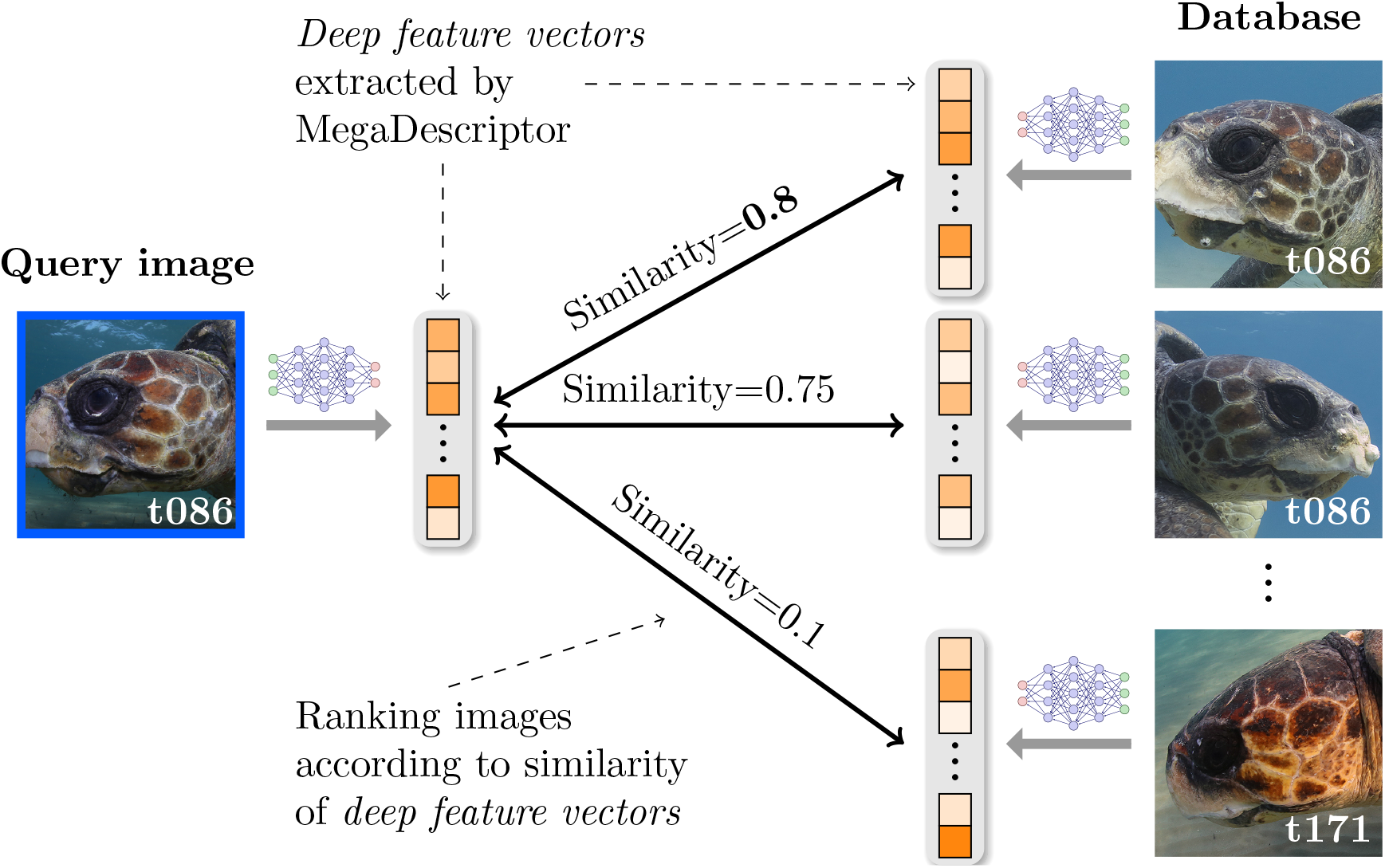
MegaDescriptor photo-ID workflow: MegaDescriptor extracts *deep feature vectors* both from the query image and the database images, which can be of any side - left or right. The database feature vectors are ranked in order of decreasing similarity (using the *cosine similarity metric*) to the query feature vectors, endowing a corresponding ranking to the database images themselves.

MegaDescriptor was trained on over 30 datasets of various species including primates, carnivores, reptiles, whales, mammals and fish. A significant subset of its training dataset is also available separately in an easily accessible form (Adam et al., 2024b). Sea turtles were included in 2 of the above 30 datasets: SeaTurtleID2022 (Adam *et al*., 2024a), of which Zakynthos-Loggerheads is an extension, as well as ZindiTurtleRecall (*T*urtle Recall: Conservation Challenge 2022) which contains green and hawksbills sea turtles ashore. Thus MegaDescriptor only saw these species in very specific selected environments. However, since it was trained on many different species, it is expected to perform well even for species unseen during its training.

### 3.3. SIFT

is a classical local feature-based algorithm which extracts a set of keypoints, i.e. locations of salient information and their corresponding descriptors (vectors describing image gradients around a keypoint) from each image (Lowe, 2004). Then the similarity score between two images is the sum of the difference between the most similar descriptors. Once similarities are computed, then the image retrieval is performed in the same way as it was described for MegaDescriptor. Since only a limited number of keypoints is extracted for each image, a single keypoint on a head side will not necessarily be selected on the other side and, therefore, SIFT-based approaches are designed to match only the same sides, as in Figure 1(a), and in general images with spatial overlap.

### 3.4. TORSOOI codes

The TORSOOI database is part a large collaborative sea turtle project which includes a web application for sea turtle photo-ID, based on TORSOOI codes. In order to generate these codes, the scales on each head profile are divided into columns and represented by a series of numbers denoting the number of edges in each scale in that column (Jean et al., 2010). The similarity between two TORSOOI codes is computed as the number of matching numbers across all columns. Therefore, it computes how many times the scales at similar locations have the same number of edges. The predictions are then made using the image retrieval setting as in the two previous methods. We note that we only applied TORSOOI to the Reunion-Greens and Reunion-Hawksbills as the codes were only available for these datasets.

The codes that we used for the implementation of the above methods is publicly available at https://github.com/sadda/sides-matching.

### 3.5. Settings for comparing sea turtle head profiles

To properly evaluate the similarities of left and right turtle profiles, we adopt five settings denoted by (A), (B), (C), (D), (E) and summarised in Table 2. Given an image, each setting defines the set of images that are to be compared with it. In the settings (A), (B) and (C) the pairwise comparisons are always done among profiles of the same individuals whereas in the settings (D) and (E), profiles of different individuals are compared. In the setting (A), images of opposite side, taken in the same year are compared. On the other hand, in the setting (B) (respectively (C)), images of the same (respectively opposite) side taken in a different year are compared. Since by dataset design, there are only four images for each individual, in each of the settings (A), (B), (C), a given image is compared with exactly one other image, see the example at the right-hand side of Table 2.

**Table 2.**
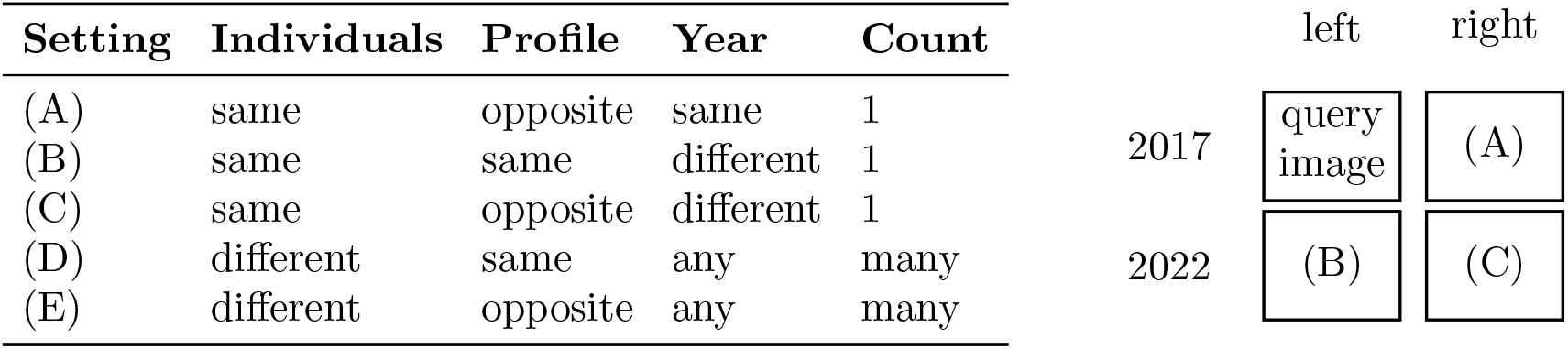
Settings for comparing sea turtle head profiles. “Individuals”, “Profile” and “Year” refer to whether images in pairwise comparisons under each setting show the same individuals or not, show the same profile or not, were taken during the same year or not. The last column “Count” refers to the number of images that each query image is compared to under each setting, see also the table at the right hand-side.

### 3.6. Evaluating the similarity of left and right profiles

In this first experiment, we compute the similarity scores given by MegaDescriptor, SIFT and TORSOOI under all settings (A)–(E) and for all the four datasets (TORSOOI for greens and hawksbills only). For example for setting (A), only similarities of pairs of images of the opposite profiles of the same individual, taken in the same year, are considered, analogously for the other settings. We recall that the average span for images taken in different years is almost four years, see Table 1. Therefore, we hypothesize that the average similarity under setting (A) will be higher than that under (C). This is because sea turtles can exhibit natural changes in skin coloration and pigmentation over the years (but also externally driven changes e.g. scratches, presence of algae), which are not side-specific. We also expect the average similarity under setting (B) to be higher than that (C) simply because of the different geometric pattern of opposite sides at a given individual. Finally, it is expected that the average similarity under (D) is roughly equal to (E) because the choice of profiles does not matter for different individuals. Note also that the deep feature vectors extracted by MegaDescriptor remain similar when the image is flipped which means that two photos of different individuals will not be judged to be less similar just because they show opposite profiles. These expectations are visualised at Figure 5.

**Figure 5.**
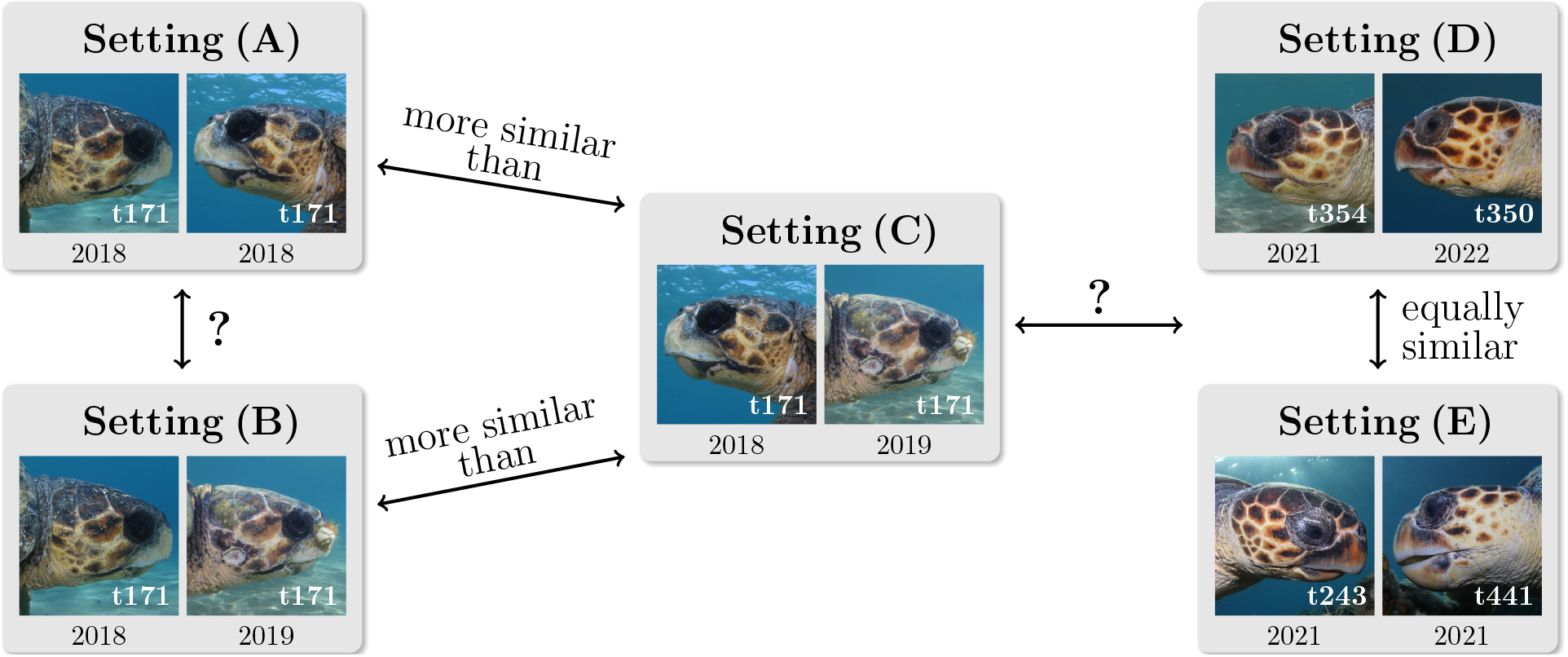
Expected relations between the average similarity of sea turtle profiles of the settings (A)–(E). The unknown relations (“?”) are closely related to the (non)-similarity of left-right profiles at a given sea turtle individual which is one of the the main investigations of this paper.

On the other hand, the relation between the average similarities under (A) and (B) is not obvious. If the former is higher, this means that it is simpler to identify a turtle from opposite profiles taken in the same year than from the same profile taken in different years. The relation between the average similarity of (C) and that of (D) is one of our main investigations. If the former is higher, it means that there is a non-negligible similarity between left and right turtle profiles of a given individual which is attributed to morphological features that are stable in time.

To numerically evaluate the differences between average similarities, we used boxplots with p-values computed by the two-sample t-test. For a comparison where we expect one mean to be larger than another (as depicted in Figure 5), we use the alternative hypothesis that one mean is larger (instead of the usual alternative hypothesis that they are different).

### Predicting sea turtle identities under the different comparison settings

To verify the results from the previous section and examine how they are translated to realistic photo-ID scenarios, we perform a series of image retrieval experiments. In particular we perform four experiments for each image in the datasets, with the latter playing the role of the query image in every experiment. For every query image, the “database” always contains all images of different individuals but we include only specific images of the same individual according to four different settings, see Table In setting (full) all the rest three images of the individual are included in the database, while settings (A)–(C) correspond to the settings from Table 2. For instance, in setting (A), the only database image that depicts the same individual to the query image, is the one of the opposite side taken during the same year, similarly for the settings (B) and (C). Furthermore, we introduce here the setting (B+C), where the database contains both sides of the individual of the query image taken in different years.

To numerically evaluate every experiment, we use the top-*k* accuracy (*k* = 1, 2, …). Here the database images are ranked in order of decreasing similarity, and if at least one of the top-*k* images shows the same individual as the query image, we deem the whole prediction as correct. The overall top-*k* accuracy for each setting is the ratio of correct predictions over the number of query images.

Top-1 accuracy reduces to the standard accuracy. The idea behind multiple ranked predictions is that the user may manually compare only a small number of images and decide whether the prediction is correct.

The experiments described in the previous section compare average similarity within each setting. Since image retrieval predictions are based on the highest similarity scores, experiments in this section compare the highest similarity withing each setting. The previous section also mentioned several relations between settings which we intend to investigate. We would like to stress that due to their design, we expect both experiments to give the same results. As an example, we expect both experiments to show that the accuracies in (A) are higher than in (C). This section also introduced the settings (B+C) and (full). The former is probably the most realistic scenario, where the query image is compared to all images from the different (previous) years. The latter is equivalent to (A+B+C) and, therefore, forms an upper bound for (A), (B) and (C). To show an example, a comparison of (A) and (full) reveals how much the performance for (A) increases by adding images from different years.

We stress again that in order to ensure that the inferred similarities in (A) are due to the turtles’s morphological characteristics, we aimed to remove as many irrelevant factors as possible: (i) We removed the backgrounds by using bounding boxes, (ii) we repeated the experiments after converting all images to grayscale to remove the effect of colorisation, and (iii) we designed the datasets so that settings (A)–(C) have the same size. The latter removes the effect of imbalanced classes and minimises the probability that a match is obtained by pure chance for individuals with many images.

## 4. Results

### 4.1. Comparison of similarities of left and right profiles

In Figure 6, we show the similarity scores for profile comparisons under all five settings (A)–(E) of Table 2, inferred by MegaDescriptor, SIFT and TORSOOI (three columns) for all four datasets (four rows). For every method, higher scores mean higher similarities, however we note that the similarity values at the horizontal axes among the different methods are not comparable. Instead we focus on the relative ordering of the average similarities of image pairs using each method under the different settings.

**Figure 6.**
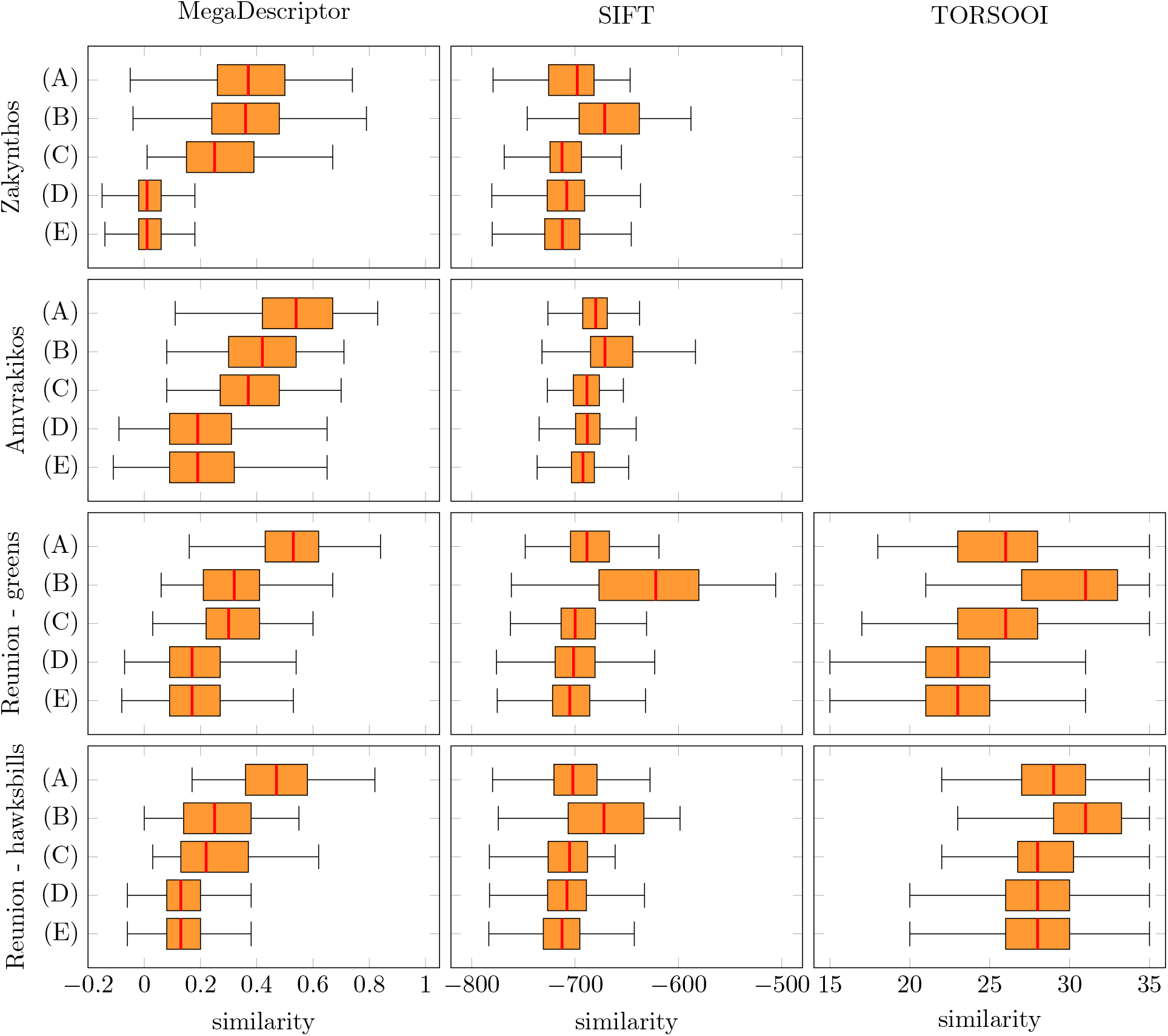
Similarity scores for profile comparisons under all five settings (A)–(E) of Table 2, inferred by MegaDescriptor, SIFT and TORSOOI (three columns) for all four datasets (four rows)

In each of the four datasets, MegaDescriptor inferred that the average similarity under (C) (same individual, opposite profile, different year) is larger than those under (D) and (E) (profiles of different individuals) with the differences being statistically significant (p-values*<* 0.001, i.e. the probability that we reject (C)>(D) is very small). This confirms quantitatively that the similarity between left and right profiles of the same individual is on average higher than the similarity between profiles of different individuals. We note that the difference between (C) and (D) & (E) is more pronounced for the Zakynthos-Loggerheads dataset, which is probably due to the fact that MegaDescriptor was trained with images of the same distribution, i.e. the SeaTurtleID2022 dataset. TORSOOI also confirmed this difference for Reunion-Greens (p-value*<* 0.001) and for the Reunion-Hawksbills (p-value=0.007). On the other hand, SIFT could not detect the similarity between left and right profiles of the same individuals, as in all datasets the similarity under (C) was not different than that under & (E) (p-values> 0.18 in all datasets). Furthermore, as we have expected, MegaDescriptor and TORSOOI do not make any distinction between the similarity under (D) (different individuals, same profile) and the one under (E) (different individuals, opposite profile), with p-values> 0.3 in all the four datasets. SIFT assigns slightly higher similarity scores in pairs under (D) compared to which suggests a dependence on image orientations for this method.

When focusing on similarities between images of the same individuals (A)–(C), we observe that MegaDescriptor infers that in all four datasets the similarity under (A) (same individual, opposite profile, same year) is at least as high as the similarity under (B) (same individual, same profile, different year). In fact in all the datasets but Zakynthos-Loggerheads there holds (A)>(B) with the difference being statistically significant (p-values*<*0.001). This difference is more evident in the Reunion-Greens and Reunion-Hawksbill datasets. On the other hand, we observe a reverse situation with SIFT, which infers that (A)*<*(B) in all datasets. This is consistent with SIFT being a local feature-based method, and as in setting (C) it assigns low similarity scores in setting (A) without being able to detect the inherit similarity of left and right profiles in an individual. TORSOOI also infers (A)*<*(B) most likely since the absolute number of scales and their edges is different in opposite profiles.

Finally as we have predicted, in general MegaDecriptor infers that the similarity under (B) is higher than that under (C). However, both settings produce relatively high similarity scores and in fact for Reunion-Greens and Reunion-Hawkbills the difference is not significant. This hints that when it comes to comparisons between photos of different years, the choice of profiles plays no significant role at least for these two latter datasets.

All the corresponding p-values for the different comparisons are collected in Table 4. There, we have highlighted with purple colour all p-values which deem the corresponding comparisons statistically significant at the level of 5%. We also present the complementary results for the grayscale images in Figure A1 and Table A1 in the Appendix. We emphasize that the relative ordering of the average similarities under the different settings remains the same as in the coloured images.

#### Examples of individuals assigned high and low left-right profile similarities

In Figure 7(top two rows), we have provided examples of photos from each dataset, of opposite profiles of the same individuals taken in the same year, i.e. setting (A), ranked with very high similarity scores according to MegaDescriptor. One can readily see the reasons that the similarity of profiles was high. For instance, looking at the first pair of the Zakynthos-Loggerheads dataset, there is high similarity due to the common whitish-texture and the not well-defined pigmentation within the scales. On the other hand, the profiles of both pairs of the Amvrakikos-Loggerheads dataset, look rather similar not only with respect to coloration but also to geometry. Focusing on the Reunion-Greens and Reunion-Hawksbills examples, we observe that while the similarity of left-right profiles is evident here as well (observe the almost identical pattern in the first Hawksbill pair), similarities on global coloration (compare the two pairs of Reunion-Greens) as well as other factors, e.g. presence of algae on both profiles at second pair of Reunion-Hawksbills, could have influenced the results. However, when looking at the pairs with the highest similarity score for the grayscale results in Figure 7(middle two rows), we do observe high similarities both in pigmentation and geometry, which makes it safe to conclude that in the absence of common global coloration, these are indeed the characteristics that lead to the assignment of high similarity scores. Related to that, in Figure 7(bottom two rows), we have also provided examples of opposite profiles of the same individuals taken in different year, i.e. setting (C), ranked with very high similarity scores. Once again one can readily notice the left-right profile similarities with respect to both coloration and geometry which are now guaranteed not to be attributed to factors other than the animals themselves.

**Figure 7.**
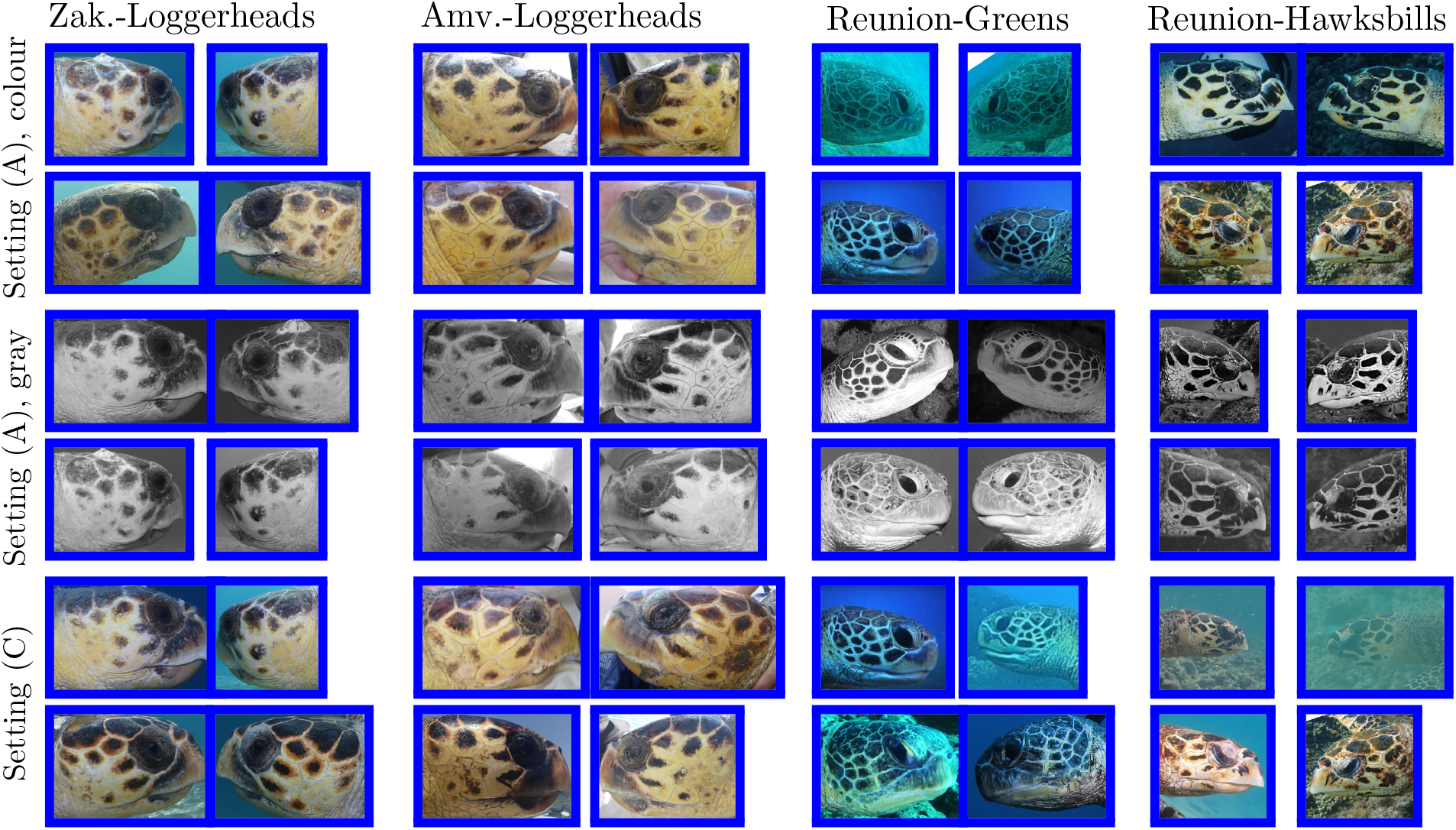
Top: Photos of opposite profiles of the same individuals taken in the same year, i.e. setting (A), ranked with the two highest similarity scores for each dataset. Middle: Corresponding photos with the two highest similarity scores but for the grayscale experiments. Bottom: Photos of opposite profiles of the same individuals taken in different year, i.e. setting (C), ranked with the two highest similarity scores for each dataset. All rankings are according to MegaDescriptor.

We can get some further insights by inspecting examples of opposite profiles of the same individuals taken in the same year, i.e. setting (A), but ranked with very low similarity scores this time, see Figure 8. For instance by visually inspecting the first pair of Zakynthos-Loggerheads, we see that the presence (resp. absence) of algae on the right (resp. left) profile could have contributed to the low similarity score. We can see further geometric and coloration differences at the Amvrakikos-Loggerheads pairs. For instance, looking at the second pair we see that the middle post-ocular scales are adjacent to one (resp. two) scale of the second scale column of the right (resp. left) profiles. We note that this pattern combination is not common for this population (KP, unpublished data). For the first green turtle pair, observe the unusual heart-shaped scale at the left profile which is absent at the right one. However, there is also a chance that the different ambient coloration could have contributed to the low similarity. Finally observe the long top-left scales at the right profiles of both Hawksbills which does not exist on the right profiles.

**Figure 8.**
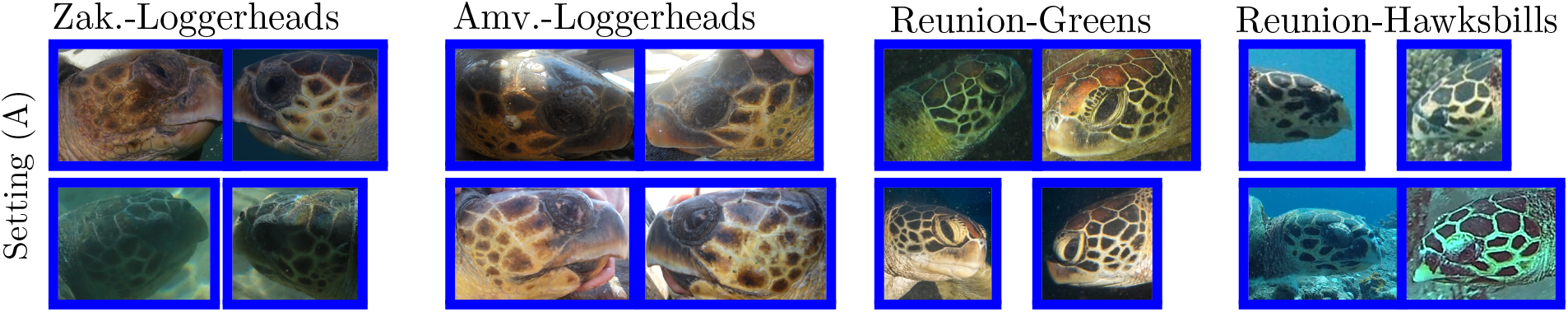
Photos of opposite profiles of the same individuals taken in the same year, i.e. setting (A), ranked with the two lowest similarity scores for each dataset, according to MegaDescriptor

### 4.2. Identity prediction under the different settings

In Figure 9, we provide the results of the series of image retrieval experiments, in terms of top-*k* accuracy for various values of *k*, under the four different settings of Table 3, for all methods and for all datasets. We also report separately the top-5 accuracies in Table 5, where for every method and dataset, we have highlighted with purple color which setting among (A), (B), (C) produced the highest top-5 accuracy.

**Table 3.**
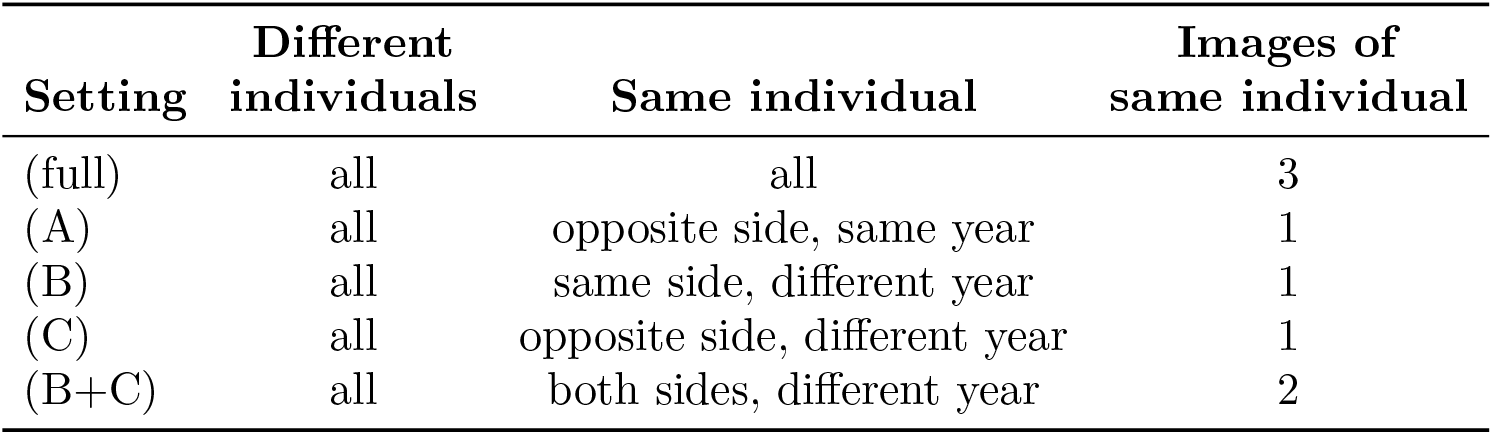
Different settings regarding the series of image retrieval experiments. “Different individuals” (respectively “Same individual”) refers to which images of the different (respectively same) individuals to the one in the query image is included in the database each time. The last column shows the number of images of the individual in the query image that are included in the database.

**Table 4.** Table of p-values computed by the two-sample t-test. The p-values that are highlighted with purple colour are the ones that deem the corresponding comparisons statistically significant at the 0.05 level.

| Method | Dataset | Alternative hypothesis |  |  |  |  |
| --- | --- | --- | --- | --- | --- | --- |
| | | (A)≠(B) | (A)>(C) | (B)>(C) | (C)>(D) | (D)≠(E) |
| MegaDescriptor | Zakynthos-Loggerheads | 0.380 | 0.000 | 0.000 | 0.000 | 0.340 |
|  | Amvrakikos-Loggerheads | 0.000 | 0.000 | 0.002 | 0.000 | 0.625 |
|  | Reunion-Greens | 0.000 | 0.000 | 0.334 | 0.000 | 0.379 |
|  | Reunion-Hawksbills | 0.000 | 0.000 | 0.239 | 0.000 | 0.464 |
| SIFT | Zakynthos-Loggerheads | 0.000 | 0.064 | 0.000 | 0.415 | 0.000 |
|  | Amvrakikos-Loggerheads | 0.000 | 0.000 | 0.000 | 0.941 | 0.000 |
|  | Reunion-Greens | 0.000 | 0.000 | 0.000 | 0.187 | 0.000 |
|  | Reunion-Hawksbills | 0.000 | 0.013 | 0.000 | 0.515 | 0.000 |
| TORSOOI | Reunion-Greens | 0.000 | 0.184 | 0.000 | 0.000 | 0.980 |
|  | Reunion-Hawksbills | 0.000 | 0.066 | 0.000 | 0.007 | 0.587 |

**Table 5.** Top-5 accuracy for the image retrieval experiments under all settings (A)–(E) of Table 3, for all methods and datasets. For every method and dataset, we have highlighted with purple color the setting among (A), (B), (C), that produced the highest top-5 accuracy.

| Method | Dataset | Setting |  |  |  |  |
| --- | --- | --- | --- | --- | --- | --- |
|  |  | (full) | (A) | (B) | (C) | (B+C) |
| MegaDescriptor | Zakynthos-Loggerheads | 0.987 | 0.919 | 0.862 | 0.694 | 0.912 |
|  | Amvrakikos-Loggerheads | 0.795 | 0.730 | 0.360 | 0.225 | 0.420 |
|  | Reunion-Greens | 0.830 | 0.765 | 0.260 | 0.255 | 0.350 |
|  | Reunion-Hawksbills | 0.882 | 0.838 | 0.390 | 0.324 | 0.463 |
| SIFT | Zakynthos-Loggerheads | 0.506 | 0.088 | 0.450 | 0.075 | 0.469 |
|  | Amvrakikos-Loggerheads | 0.415 | 0.070 | 0.370 | 0.020 | 0.375 |
|  | Reunion-Greens | 0.680 | 0.085 | 0.650 | 0.010 | 0.655 |
|  | Reunion-Hawksbills | 0.559 | 0.118 | 0.507 | 0.037 | 0.515 |
| TORSOOI | Reunion-Greens | 0.780 | 0.315 | 0.720 | 0.265 | 0.765 |
|  | Reunion-Hawksbills | 0.618 | 0.184 | 0.507 | 0.140 | 0.559 |

**Figure 9.**
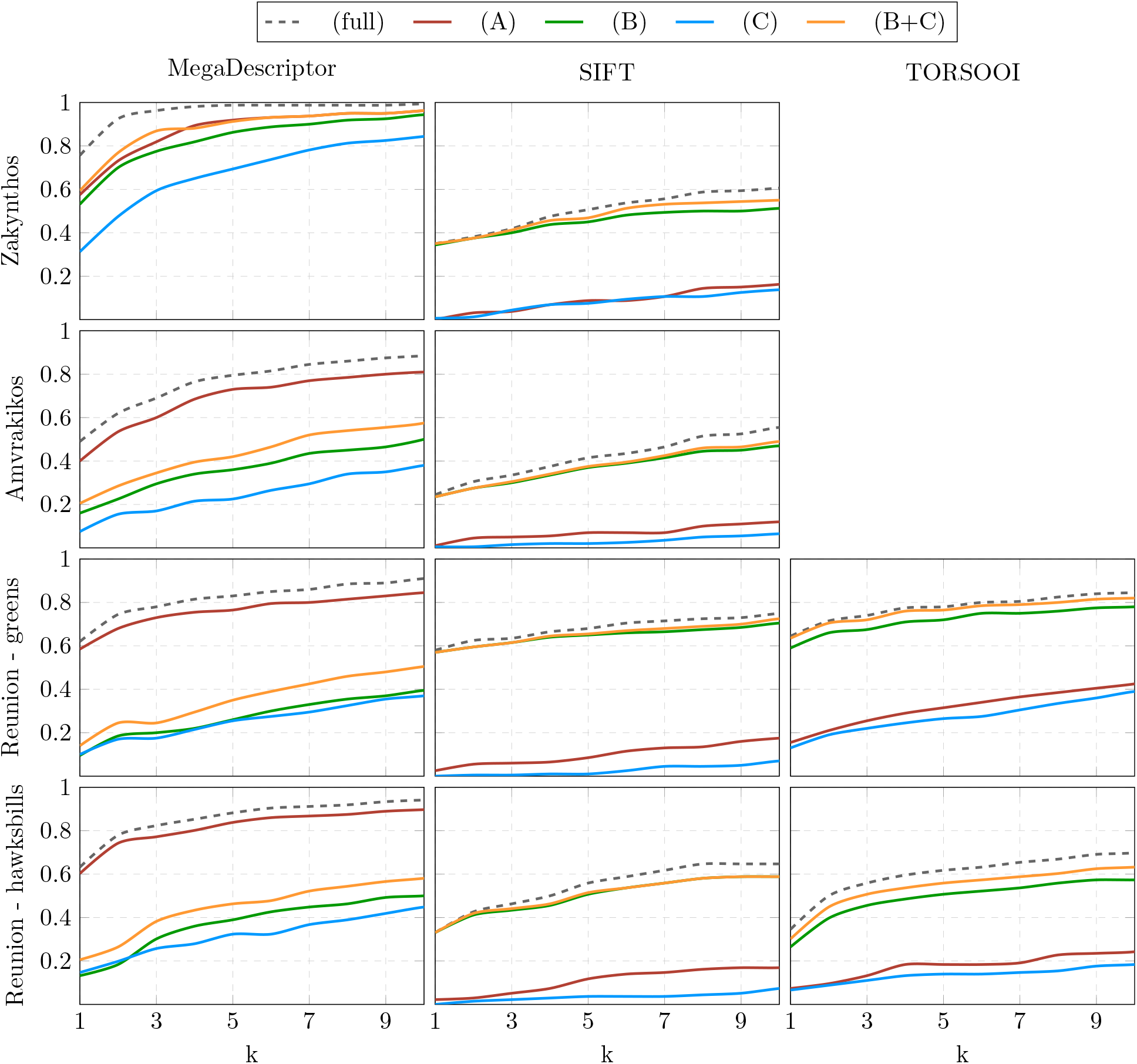
Top-*k* accuracies for the series of image retrieval experiments, for various values of *k*, under all settings (A)–(E) of Table 3, for all methods and for all datasets

Focusing first on the Zakynthos-Loggerheads dataset, which is the one that MegaDescriptor is better trained for, we observe much higher accuracies than SIFT under all settings, compare top two plots in Figure 9. For instance, with regards to the - “classical” for sea turtle photo-ID - setting (B) where one looks to match the query image with one of the same profile taken in a previous year, MegaDescriptor has a top-5 accuracy of 86.2% versus 45.0% for SIFT, compare also the green lines in the two aforementioned plots. Notably, under the most challenging setting (C), where the only database image of the query individual is one of the opposite profile taken during a different year, MegaDescriptor achieved an accuracy of 69.4% vs only 7.5% for SIFT, compare also the blue lines. The low accuracy of SIFT with respect to the latter setting is expected since, as the previous experiment confirmed, it cannot detect similarities of left and right profiles. Under the setting (B+C), i.e. when including in the database both profiles of the individuals, both taken in a different year the MegaDescriptor accuracy rises to 91.2%. This is comparable with the accuracy under setting (A) which is 91.9%.

In fact, MegaDescriptor’s accuracy under (A) was the highest among the settings (A)–(C) for all datasets, see for instance the values highlighted in purple in Table 5. MegaDescriptor’s accuracy under (A) is also much higher than the other methods under this setting for all datasets, resulting to similarly higher accuracy when the (full) setting is considered, see the “(full)” column in Table 5 and the dotted lines in Figure 9.

The accuracy of MegaDescriptor under (B) and (C) was not as high in the other three datasets, in fact both SIFT and TORSOOI performed significantly better in Reunion-Greens and Reunion-Hawksbills under setting (B), 65.0% & 72.0% vs 26% and 50.7% & 50.7% vs 39.0% respectively. Notably, particularly in the Reunion datasets, the accuracy of MegaDescriptor under (C) was only slightly lower to that under (B), 25.5% vs 26.0% and 32.4% vs 39.0% for Reunion-Greens and Reunion-Hawksbills respectively. This is consistent with the results of Figure 6 regarding the approximate equality of settings (B) and (C). The difference was more pronounced for Amvrakikos-Loggerheads, 22.5% vs 36.0%. This again indicates that when ones aims to match a query image of a turtle to a database of turtle images taken in previous years, the choice of profile may play little role, compare green and blue lines in the first column of Figure 9.

## 5. Discussion

Sea turtle photo-ID has seen a remarkable progress over the years with respect to both expanding its range of ecological applications as well as the ever increasing improvement of automated techniques. Up to now, all the current widely used automated photo-ID methods, (Dunbar *et al*., 2014; Calmanovici *et al*., 2018; Jean *et al*., 2010; Dunbar *et al*., 2021; Mills *et al*., 2023), as well those based on manual divide-and-conquer strategies (Schofield *et al*., 2008; Papafitsoros *et al*., 2024), are local feature-based or species-specific methods operating under an side-specific image retrieval setting where only comparisons between the same profiles are performed, i.e. left vs left or right vs right. Our results suggest that MegaDescriptor and in general the ever improving deep feature-based methods (Oquab *et al*., 2024; Cermak *et al*., 2024; Nepovinnykh *et al*., 2024) will result to a change of paradigm in sea turtle photo-ID due to their ability to assign high similarity scores not only to pairs of the same profiles of the same individual but also to pairs of opposite profiles. Here for the first time, we showed quantitatively (i) an inherit “deep” similarity of left and right profiles of a given individual, which was on average significantly higher than those of different individuals, and (ii) we demonstrated how this similarity can be translated to improved accuracies of these methods under image retrieval experimental settings which mimic realistic sea turtle photo-ID matching processes.

The higher similarity of left and right profiles of a given individual compared to the one between different individuals was mainly detected by MegaDescriptor, and partly by TORSOOI. However, this similarity was completely unseen by SIFT, whose extracted keypoints contain only local information, i.e. gradients contained in a circle of some radius, which is unlikely to occur in the same spatial position on both sides. On the other hand, the deep feature vectors of MegaDescriptor, encapsulate also global information of the two sides which are more common in the same individual. The similarity was mainly manifested in the relationship (C)>(D) which was significant for all four datasets. The fact that compared images in (C) were taken in different years diminished the danger of image pairs being assigned with high similarity scores due to factors other than the turtles’ characteristics like common background or similar global coloration. However, we could not quantify to what degree this assigned high similarity was due to the similar geometrical patterns of the scales, the inherit coloration and pigmentation of the individual or other characteristics e.g. shape of the head (Casale *et al*., 2017). Nevertheless, the relationship (C)>(D) also held for the grayscale version of the experiments, hinting that features other than colouration did indeed influence the similarity scores. Coloration, pigmentation and texture is common throughout the head of an individual turtle and as such, high similarity of the left and right profiles with respect to those is not surprising. The reasons that skin coloration and pigmentation are different among individuals are not so well understood but are likely attributed to a combination of genetic factors, differences in foraging habits and sun exposure (Papafitsoros *et al*., 2024). On the other hand, there are essentially no theories that could explain the reasons behind the geometric similarity between left and right profiles. Thus, further research is required in particular with respect to the mechanisms behind scale formation at the embryonic stages (Moustakas-Verho et al., 2014; Zimm, 2019).

While geometrical scale patterns are stable throughout a turtle’s life span (Carpentier *et al*., 2016), pigmentation and coloration on a turtle’s facial skin can change due to various factors like aging, shifts of foraging habits, and seasonal changes in sun exposure (Adam *et al*., 2024a; Papafitsoros *et al*., 2024). Factors like presence of algae, injuries and scratches can also change the appearance of a turtle year after year. This could explain the high similarity scores that were assigned by MegaDescriptor to opposite pairs of the same individual taken in the same year (setting (A)) compared to pairs taken in different years (setting (C)). In fact, it turned out that for MegaDescriptor it was easier to recognize individuals based on comparison of photos of opposite profiles of an individual taken in the same year, compared to comparisons of the same profiles taken in different years (setting (B)). We could not entirely exclude the possibility that common backgrounds and global coloration, which are often similar in photos taken during the same encounter (Adam et al., 2024a; Adam et al., 2024b), could have artificialy inflated the similarities in (A) despite our efforts to mitigate this effect by converting images to grayscale. It is still possible that other factors that could not be eliminated by this conversion, such as same degree of focus or use of strobe lights on photos of same encounter resulting in increased contrast and scale definition, could have have influenced these results. This could be the case for the Reunion-Greens and Reunion-Hawksbills datasets where the relationship (A)>(B) was more pronounced. On the other hand, we note that (A)>(B) holds, albeit to a lesser degree, also for the Zakynthos-Loggerheads and Amvrakikos-Loggerheads datasets despite the fact that the photo-capturing conditions where essentially uniform across the years. Thus we argue that the high similarity scores in (A) compared to (C), can be indeed attributed to changes in the turtles’ appearance.

The detection of left-right profile similarities by the MegaDescriptor has several positive implications for sea turtle photo-ID. We note first that with regards to the image retrieval experiments, it was evident that MegaDescriptor’s accuracy for the Zakynthos-Loggerheads dataset was much higher than the other three datasets. In fact, MegaDescriptor’s performance on this dataset was such that it could retrieve the query individual even when only the opposite profile taken in a different year was available (setting (C)) with an almost 70% top-5 accuracy. We attribute this to the fact that images of individuals from the SeaTurtleID2022 dataset were part of the training set of MegaDescriptor. Even though the individuals of Zakynthos-Loggerheads were not part of SeaTurtleID2022, turtles from both of these datasets belong to the same population, thus sharing similar morphological characteristics and their photos were taken under similar conditions, with similar camera and by the same photographer. It is known that deep feature and in general neural network-based methods become more accurate when they are trained on images of the same characteristics (same distribution) as the ones they are tested on (Goodfellow *et al*., 2016). Thus we expect that the performance of MegaDescriptor on other datasets can be significantly improved by simply fine-tuning its parameters using images from these datasets. Nevertheless, despite the decreased accuracy, we generally observed in all datasets that when comparing photos of different years, opting to base the comparison on the same profiles provides only slight advantage over doing it so based on the opposite profiles and in fact there is a clear gain in using both. It is particularly useful to be able to perform sea turtle photo-ID under a non side-specific setting, as data collection protocols e.g. citizen scientist submissions, camera traps or skittish animal behaviour can often result to obtaining only one side of the animal’s head. Being unable to match opposite sides can pose certain obstacles when using photo-ID in ecological applications, and it requires careful statistical analysis (McClintock et al., 2013).

The benefits of allowing comparisons of opposite profiles when attempting to match a single query image, emphasizes the superiority of profile-based sea turtle photo-ID in comparison to photo-ID based on the turtle’s dorsal scale patterns. It is known that the more photos of an individual already exist in the database, the higher the chances that a query photo of that individual will be matched to at least one of those, resulting to a correct identity prediction (Dunbar *et al*., 2014). In our experiments, that was evident for instance in the increased accuracies under settings (full) and (B+C) in comparison to settings (B) and (C). Thus simply due to the fact that there are two profiles per individual and only dorsal side, the database can be twice as rich. However we should note that dorsal-based sea turtle photo-ID is still the only option when drones are the main means for capturing photos (Comis et al., 2022).

Our work further highlighted the need for multiple well-curated, publicly available multiple year-spanned sea turtle datasets that can be used both for development of algorithms (training) and their proper evaluation (testing). In particular it is crucial that timestamps i.e. capture dates, as well as orientation labels of the heads, are included in the metadata of each image. For instance, training deep feature-based methods with photos of individuals spanning several years, could force the extracted feature vectors to encode exactly these identifying characteristics that are stable in time. This is particularly crucial for developing methods that can identify individuals over large time scales, e.g. from juveniles to adults. Maturation of sea turtles up to a point where they stop growing can take decades (Baldi et al., 2023), making the assemblage of such datasets particularly challenging. On the other hand, adding timestamps in each photo is simple yet crucial for proper method evaluation (Adam et al., 2024a). By comparing the query images only with those taken in different years, allows to mimic realistic photo-ID matching workflows and avoids the comparison of photos taken in the same conditions/encounters which could artificially inflate the accuracy of algorithms due to the common background or global coloration. We also encourage the use of multiple datasets, with multiple species, created with different settings. The majority of sea turtle photo-ID studies typically use a single dataset for a single location to perform experiments. However here are showed how the results of a method can vary on different datasets and how this can be explained based on their characteristics. At the moment there are only two publicly available sea turtle datasets for algorithmic training and testing SeaTurtleID2022 (Adam *et al*., 2024a), and ZindiTurtleRecall (*T*urtle Recall: Conservation Challenge 2022), and we strongly recommend researchers to publish also their own datasets. Here we contribute to that need by making all our four datasets publicly available.

## 6. Conclusion

In this work we showed that the higher-within-individuals similarity of left and right profiles in sea turtles can be detected by state-of-the-art deep feature re-identification methods with clear practical benefits. While our current work focuses on sea turtles, it paves the road for exploiting morphological symmetries of opposite sides of animals when performing photo-ID with deep neural networks. Such detectable symmetry for instance has been shown to exist in dolphins (Genov *et al*., 2018) and it is likely to exist in other species whose identification has been so far performed under side-specific settings or in general based on image pairs of some spatial overlap. We thus anticipate further research towards this direction which will be facilitated by the constant progress of computer vision and artificial intelligence on one hand and the ever increasing availability of public databases of wild animals (Čermák et al., 2024; Shinoda and Shiohara, 2024).

## Code and Data availability

Code is available at:

https://github.com/sadda/sides-matching.

Datasets are available at:

https://www.kaggle.com/datasets/wildlifedatasets/zakynthosturtles

https://www.kaggle.com/datasets/wildlifedatasets/amvrakikosturtles

https://www.kaggle.com/datasets/wildlifedatasets/reunionturtles

## Funding

This research has been supported by the Ministry of Education, Youth and Sports of the Czech Republic under project SGS-2024-017.

## CRediT authorship contribution statement

**Lukáš Adam:** Code development, experiment design & execution, data curation, writing original draft. **Kostas Papafitsoros:** Conceptualization, experiment design, data provision & curation, writing original draft. **Claire Jean:** Data provision & curation, manuscript review. **ALan Rees:** Data provision & curation, manuscript review.

## Appendix A. Experiments on grayscale images

In this appendix section, we provide results on grayscale images which correspond to the same experimental settings as for coloured images presented in the main manuscript body. Thus, Figures A1 and A2 correspond to Figures 6 and 9, respectively, while Tables A1 and A2 correspond to Tables 4 and 5, respectively.

**Figure A1.**
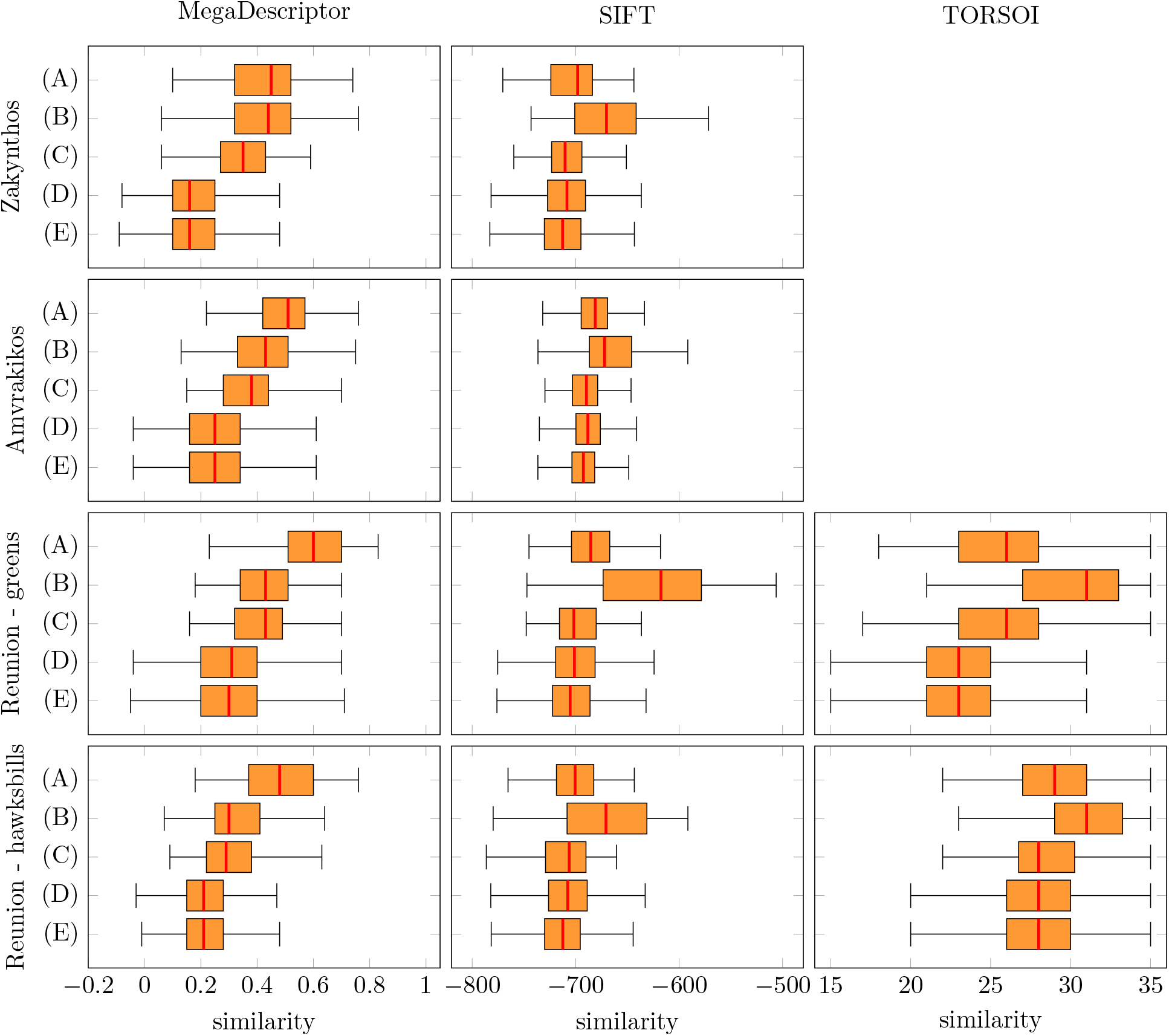
Grayscale images: Similarity scores for profile comparisons under all five settings (A)–(E) of Table 2, inferred by MegaDescriptor, SIFT and TORSOOI (three columns) for all four datasets (four rows)

**Table A1.** Grayscale images: Table of p-values computed by the two-sample t-test. The p-values that are highlighted with purple colour are the ones that deem the corresponding comparisons statistically significant at the 0.05 level.

| Method | Dataset | Alternative hypothesis |  |  |  |  |
| --- | --- | --- | --- | --- | --- | --- |
|  |  | (A)≠(B) | (A)>(C) | (B)>(C) | (C)>(D) | (D)≠(E) |
| MegaDescriptor | Zakynthos-Loggerheads | 0.855 | 0.000 | 0.000 | 0.000 | 0.212 |
|  | Amvrakikos-Loggerheads | 0.000 | 0.000 | 0.000 | 0.000 | 0.505 |
|  | Reunion-Greens | 0.000 | 0.000 | 0.067 | 0.000 | 0.434 |
|  | Reunion-Hawksbills | 0.000 | 0.000 | 0.042 | 0.000 | 0.681 |
| SIFT | Zakynthos-Loggerheads | 0.000 | 0.176 | 0.000 | 0.206 | 0.000 |
|  | Amvrakikos-Loggerheads | 0.000 | 0.000 | 0.000 | 0.946 | 0.000 |
|  | Reunion-Greens | 0.000 | 0.000 | 0.000 | 0.202 | 0.000 |
|  | Reunion-Hawksbills | 0.000 | 0.004 | 0.000 | 0.722 | 0.000 |
| TORSOOI | Reunion-Greens | 0.000 | 0.184 | 0.000 | 0.000 | 0.980 |
|  | Reunion-Hawksbills | 0.000 | 0.066 | 0.000 | 0.007 | 0.587 |

**Table A2.** Grayscale images: Top-5 accuracy for the image retrieval experiments under all settings (A)–(E) of Table 3, for all methods and datasets. For every method and dataset, we have highlighted with purple color the setting among (A), (B), (C), that produced the highest top-5 accuracy.

| Method | Dataset | Setting |  |  |  |  |
| --- | --- | --- | --- | --- | --- | --- |
|  |  | (full) | (A) | (B) | (C) | (B+C) |
| MegaDescriptor | Zakynthos-Loggerheads | 0.906 | 0.669 | 0.612 | 0.437 | 0.719 |
|  | Amvrakikos-Loggerheads | 0.740 | 0.595 | 0.365 | 0.170 | 0.405 |
|  | Reunion-Greens | 0.730 | 0.665 | 0.250 | 0.185 | 0.300 |
|  | Reunion-Hawksbills | 0.838 | 0.750 | 0.324 | 0.279 | 0.434 |
| SIFT | Zakynthos-Loggerheads | 0.500 | 0.081 | 0.444 | 0.062 | 0.469 |
|  | Amvrakikos-Loggerheads | 0.405 | 0.050 | 0.365 | 0.025 | 0.370 |
|  | Reunion-Greens | 0.660 | 0.105 | 0.625 | 0.030 | 0.630 |
|  | Reunion-Hawksbills | 0.537 | 0.088 | 0.485 | 0.029 | 0.493 |
| TORSOOI | Reunion-Greens | 0.780 | 0.315 | 0.720 | 0.265 | 0.765 |
|  | Reunion-Hawksbills | 0.618 | 0.184 | 0.507 | 0.140 | 0.559 |

**Figure A2.**
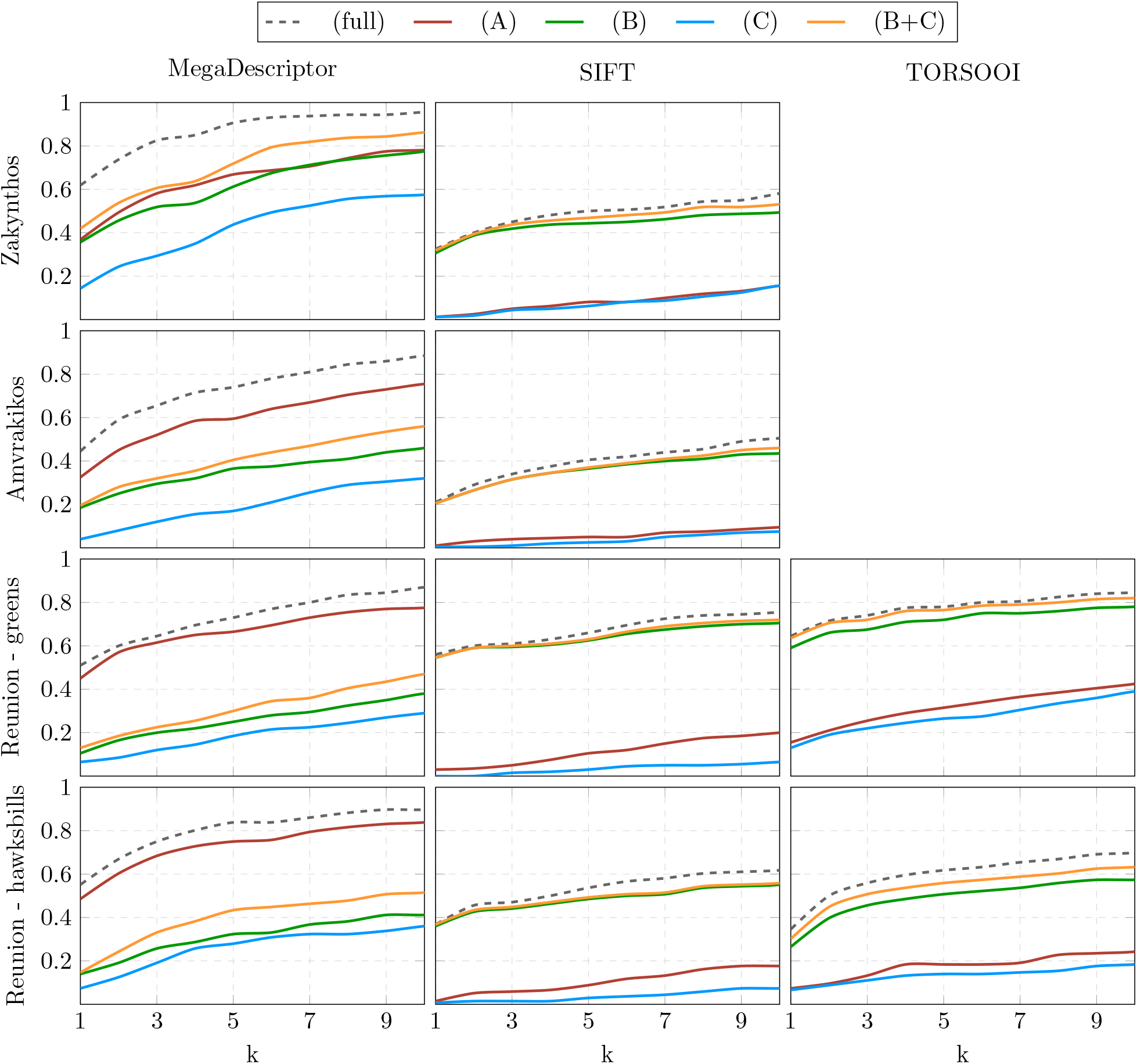
Grayscale images: Top-*k* accuracies for the series of image retrieval experiments, for various values of *k*, under all settings (A)–(E) of Table 3, for all methods and for all datasets

## Notes

### Competing Interest Statement

The authors have declared no competing interest.

